# Structural genomic variations and their effects on phenotypes in *Populus*

**DOI:** 10.1101/2023.02.14.528455

**Authors:** Ilga Porth, Roos Goessen, Berthold Heinze

**Affiliations:** Département des sciences du bois et de la forêt, Université Laval, 1030 Ave de la Médecine, G1V 0A6 Québec City, Canada; Federal Research and Training Centre for Forests, Natural Hazards and Landscape, Seckendorff-Gudent-Weg 8, 1131 Wien, Austria

**Keywords:** adaptation, copy number variation, deletions, gene dosage, pangenome, quantitative traits

## Abstract

DNA copy numbers have recently emerged as an important new marker system. In the absence of a contiguous reference genome, alternative detection systems such as the comparative hybridization method have been used to detect copy number variations (CNVs). With the advent of chromosome-level resolved reference genomes based on the incorporation of long-read sequencing and powerful bioinformatics pipelines, comprehensive detection of all structural variations (SVs) in the poplar genome is now within reach. Gene CNVs and their inheritance are important because they can cause dosage effects in phenotypic variations. These are potent genetic markers that should be considered in complex trait variation such as growth and adaptation in poplar. SVs such as CNVs could be used in future genomic selection studies for poplar, especially in cases when heterosis increases hybrid performance (hybrid vigor). This Chapter reports recent findings on SVs in natural populations of *Populus* spp. as well as on artificially induced SVs in poplar to understand their potential importance in generating a considerable amount of phenotypic improvement. The Chapter concludes with an outlook on the future implementation of knowledge on SVs in poplar crop breeding.

## 2.1 Introduction

Structural variations (SVs) are distinguished from small genetic variation (single nucleotide polymorphisms (SNPs); short insertion/deletions (indels)) found within genomes by sequence blocks of at least 50 nucleotide base pairs (bp) in length (Ho et al., 2020), and are classified into indels, inversions, duplications, and translocations (Sedlazeck et al., 2018a). SVs can even reach impressive dimensions in the scale of megabases (Ho et al., 2020). SVs can be found within or between chromosomes; they are identified between individuals of the same species or different species within a single lineage. SVs are also an important source of genetic information when it comes to causes of human diseases (Sedlazeck et al., 2018a). Copy number variants (CNVs) represent a subclass of SVs that are larger stretches of DNA sequence information, typically greater than 1,000 bp and with sequence identity above 90%, that may be found duplicated, inserted, or deleted in the genome, or referred to as presence/absence variants (PAVs). CNVs can also be found in DNA segments that cover genes and are accordingly termed gene copy number variations.

Knowledge about the presence of gene duplications in eukaryotes dates back to the 1930s. In fact, karyotypically aberrant and microscopically visible SVs have been described early on for non-lethal human disease (*e*.*g*., Down Syndrome: Trisomy 21). The more systematic identification of SVs commenced in the early 2000s with the availability of advanced microarray technology mainly driven by human medical research. Thereby, it was found that SVs in humans were common rather than rare events, and newest estimates report c.20,000-25,000, depending on the discovery approach (Escaramis et al., 2015; Sedlazeck et al., 2018a; Ho et al., 2020). Generally speaking, different genomics technologies can discover different types of SVs. High-throughput whole genome sequencing based on short reads (Illumina, second-generation sequencing technology) discovers most effectively inversions or translocations (*i*.*e*., balanced SVs) and typically smaller SVs as well as novel insertions (Ho et al., 2020), since this application does not rely on a reference, whereas comparative genomic hybridisation (array technology) requires probe design and is more effective in detecting indels and PAVs (unbalanced SVs, CNVs). Nevertheless, short-read sequencing is prone to high error rates when it comes to SVs discovery (Sedlazeck et al., 2018a) and is ineffective in resolving inter- and intra-chromosomal translocations, especially large (greater 30,000 bp) ones (Ho et al., 2020). To overcome the limitations of these two technologies, third generation sequencing (PacBio^®^, ONT^®^) as well as scaffolding techniques (Hi-C, optical mapping) have proven effective in resolving complex structures (Sedlazeck et al., 2018a; Ho et al., 2020; Canaguier et al., 2022). However, these technologies lack the base calling accuracy information provided by short reads. Overall, only the use of a multiplatform approach that combines the individual advantages of each SVs detection approach will ultimately enable comprehensive detection of all SVs in a genome (Ho et al., 2020). However, this may be prohibitively expensive for the majority of individual research laboratories. Recently, a new approach to identify notoriously difficult-to-assess translocations has been proposed that does not require genotyping and instead uses dosage of sequence reads binned into genomic segments defined by sequence coverage along the reference genome. This method works best in segregating populations where linkage disequilibrium between copy number variants is assessed (Comai et al., 2021).

The mechanisms underlying the formation of SVs are non-allelic homologous recombination and replication-based mechanisms; they depend on local repeat structures (such as the presence of low-copy repeats), and they result in breakpoints within these repetitive regions (Ho et al., 2020). Bioinformatics can then exploit signatures resulting from the mapping discrepancies between the read from a sample and the reference genome, whereby read-pair bioinformatic approaches evaluate the alignment orientation and the distance of paired ends, while the read-depth approaches detect deletions or duplications due to the divergences in mapping depth, and so-called split-read methods exploit the alignments mapping over SV breakpoints to detect translocations or inversions (*ibidem*).

Transposable elements (TEs) are considered at the base of shaping many structurally relevant aspects in eukaryotic genomes that involve chromosomal rearrangements including the formation of SVs; TE-rich genomic regions are known to evolve rapidly as are SVs in comparison to SNPs. TEs also shape transcription factor-binding sites; due to their differential expansion in the diploid progenitor genomes, TEs in combination with DNA methylation patterns can contribute to the transcriptional plasticity in allopolyploids. In diploid plant species, the main source of structural variation related to the traditional PAVs and CNVs is due to the involvement of TEs. Genomic SVs are generally recognized as an important determinant of agricultural traits (e.g., yield-related, adaptation), therefore, SVs have sparked increased interest for breeding targets (Schiessl et al., 2019). In fact, the genetic improvement of widely used polyploid crops (such as rapeseed or wheat) is tightly linked to such SVs (Schiessl et al., 2019). In addition to traditional PAVs and CNVs, allopolyploid species that arose from the combination of two (or more) different genomes as opposed to autopolyploids generated from single genome duplication, show homeologous genomic recombinations during meiosis where genetic material between the different subgenomes originating from the (two) *ancestrally* related chromosomes is exchanged. Hence, owing to the nature of their generation, allopolyploid species show more complexity in genomic SVs and these also include different translocation events for homeologous genetic material (Schiessl et al., 2019). The complexity and size of the genome poses a bioinformatics challenge when it comes to accurately decoding the genome of a polyploid species, identify the effect of the homeologous genetic exchanges on phenotypes, including gene expression, but also for determining allelic SNPs in polyploids. Therefore, also in view of higher accuracy in determining SVs in polyploid species, a hybrid sequencing approach involving third-generation sequencing technologies providing now increasingly accurate long reads (10,000-30,000 bp on average) from ultra-long DNA molecules is advisable (Schiessl et al., 2019; Ho et al., 2020).

An important source of genomic variation within a species is unravelled as presence or absence of genes in sequenced individuals. More specifically, this refers to the pangenome concept, which generally describes the sum of the (stable) core genome, i.e., genes which are present in all genomes of a species, and the (variable) accessory genome of an entire species. Unbalanced SVs (CNVs; PAVs) are referred to as the dispensable genome (Marroni et al., 2014). Interestingly, plant genes associated with biotic and abiotic stresses have been found to be frequently enriched in the variable (dispensable) genes group (Bayer et al., 2020). Knowledge about the pangenome would help better understand the functionality of genomes at large. A high-quality pangenome reference is also necessary to improve the mappability of resequencing data from germplasm collections and thus improve our capability to capture important natural variation in a species (Wendel et al., 2016). Current methodologies to deduce the pangenome are related to whole genome comparison, PAVs calling, or general patterns of genomic diversity and conservation depending on the computational approach used (Bayer et al., 2020). The elucidation of pangenomes in plants is certainly more complex than in bacteria for which the presence of a pangenome was first described in 2005 (*Streptococcus*; Tettelin et al., 2005), and high quality long-read sequencing is at the base of finding such SVs in plant genomes with high confidence. Studies in plants that have identified gene PAVs have increasingly become available only since 2014 (for a first compilation of 20 of such studies see Bayer et al., 2020). More specifically, those studies defined the core genome in percentage of genes or gene clusters that are common to all accessions of a species, in addition to the ‘dispensable’ genes or gene clusters that are absent from the reference. However, these studies utilized a highly variable number of accessions (3-3,000), rendering information on the respective core genomes also highly variable (39-92%). Intuitively, the higher the number of accessions (isolates) sequenced for a species, the higher the uncovered extent of variable genome content (higher number of pangenome genes) as already shown in *Streptococcus*. However, for those plant studies mentioned in Bayer et al. (2020), a dependency between the utilized sample size and the uncovered core genome extent in a species could not be deduced, probably due to the phylogenetic diversity in the samples. An important finding from those studies was also that there are subpopulation-specific core genes, while the same genes appear to be variable across subpopulations, suggesting that such SVs (especially PAVs) contribute to population substructure in a species (see also Stölting et al., 2015, for CNVs of the *UDP-glucuronate 4-epimerase 3* gene in populations of *Populus alba*). TEs may contribute to gene variability (gene loss or gene birth), and *de novo* genes might have their origins in long non-coding transcripts. Variable genes content is also increasingly considered in genetic association analyses with agronomic traits of interest (yield; disease resistance). Plant domestication has in fact contributed to a reduction in genomic diversity, often accompanied by gene loss, as suggested by recent plant pangenome studies. Breeding can (re-)introduce genes of interest, including those shown to be important for adaptation to the environment.

In forest trees, there have been few studies on genomic SVs. There are several reasons for this: First, tree genomes are among the most complex plant genomes, especially those of conifers, lacking contiguous genome assemblies; second, the recognition that SVs are important for economically and ecologically important trait variation and would therefore be relevant for tree breeding has lagged behind that of agricultural crops; third, approaches to study SVs in trees are mostly limited to array technology of comparative genomic hybridisation (aCGH), and focused on gene space, but are less inclusive of whole-genome sequencing data at the moment. While the first hints of copy number variation in conifers came from a few earlier studies (Harry et al., 1989; Kinlaw and Neale 1997; Kvarnheden et al., 1998), the first targeted CNV studies in a conifer were conducted by John MacKay’s research group at Laval University using aCGH technology. Their two studies (Prunier et al., 2017a; Prunier et al., 2017b) showed that gene CNVs are an important source of adaptive evolution in spruce. Natural poplar accessions were first screened for genome-wide SVs by Pinosio et al. in 2016, and as early as 2015, Andrew Groover’s research group at UC Davis artificially induced SVs by pollen irradiation, generating a large set of indel mutations in a hybrid poplar progeny (Henry et al., 2015), which were used to study the effects of gene dosage on various quantitative traits in poplar. As can be seen, the evaluation of SVs in poplar is very new and served several purposes: to evaluate naturally occurring SVs in poplar, (2) to study induced SVs in poplar. The following sections of this Chapter provide an overview of recent advances in SV research in poplars, with an emphasis on linking genotype and phenotype.

## 2.2 Naturally occurring SVs in poplar

The first pangenome in poplar obtained through genome re-sequencing was published in 2016. Pinosio et al. (2016) used Illumina DNA short reads, obtained at an average sequencing depth of 5x, from a limited number of 22 individuals (19 of which were *Populus nigra*) to infer indel SV features among three poplar species and inferred the interspecific core genome with 80.7% shared genes; most of the dispensable poplar genome (indels) was attributed to species-specific variation. As the authors noted, the higher proportion of the dispensable genome (indels) unique to *P. nigra* could be due to the fact that this species was represented by more individuals in their sample set compared to *P. deltoides* and *P. trichocarpa*. If indels were found, they were mostly located outside of the gene coding regions likely due to purifying selection, and these SVs were also kept at low expression levels or not expressed at all (pseudogenes). Genes directly affected by SVs in their coding region exhibited higher rates of non-synonymous base changes, indicative of fast evolving gene products if expressed. Pinosio et al. also evaluated gene CNVs for five high coverage individuals using the depth of coverage signature for all reads uniquely aligning on each of the 41,335 gene features of the annotated reference Nisqually-1 (*P. trichocarpa*) and found that, among all the SVs features associated with genes, CNVs were overrepresented among SV features for which genes were least expressed. An overrepresentation of CNVs related to disease resistance genes was noted as well. Overall, c.8% of all gene features in poplar showed genic CNVs, however, the distribution of these SVs was not random, as gene CNVs appeared preferentially near telomers, close to TEs, and tended to cluster in gene-rich regions on chromosomes. Inter-specific variation was generally higher than intra-specific variation for all SV features studied.

Another recent study specifically investigated the occurrence of intra-specific gene CNVs in a widely distributed poplar species in North America. Using aCGH technology, Prunier et al. (2019) screened 19,823 gene sequences which presented a subset of the originally targeted 41,337 known poplar genes for CNVs, thereby testing two intra- and inter-provenance full-sib crosses created for *Populus balsamifera*. The authors used multiple probe quality criteria outlined in the article such as a minimum probe number of five per gene, a minimum DNA segment length queried by probes, and required location of genes within the 19 large chromosomic scaffolds of the *P. trichocarpa* reference genome. They further designed a custom 4×180k probe array with on average 8.5 probes per gene and employed a hybridisation design adapted to maximize the number of subarrays available for between-individuals’ hybridisations, while also accounting for interarray variability. A higher number of gene CNVs was detected for the inter-provenance population, which is in line with its greater genetic diversity expected on the basis of the higher parental genetic distance for this progeny. The results for balsam poplar showed that copy losses in gene CNVs were more prevalent. Moreover, the majority (68%) of gene CNVs was found located over several CNV hotspots, mainly near telomeric regions (**Figure 2.1**). Rare *de novo* CNVs also clustered in such CNV hotspots, a strong indication that those genomic regions are highly dynamic in terms of creating genetic novelty. Because Prunier’s work in balsam poplar also found an important overlap with the CNV hotspots of *P. nigra* and *P. deltoides* (83%) discovered in Pinosio’s work (**Figure 2.1**), a large number of hotspots are most likely shared across the genus of *Populus* and probably beyond. This work also identified adaptive gene CNVs, further discussed in subsequent sections of this Chapter.

**Figure 2.1.**
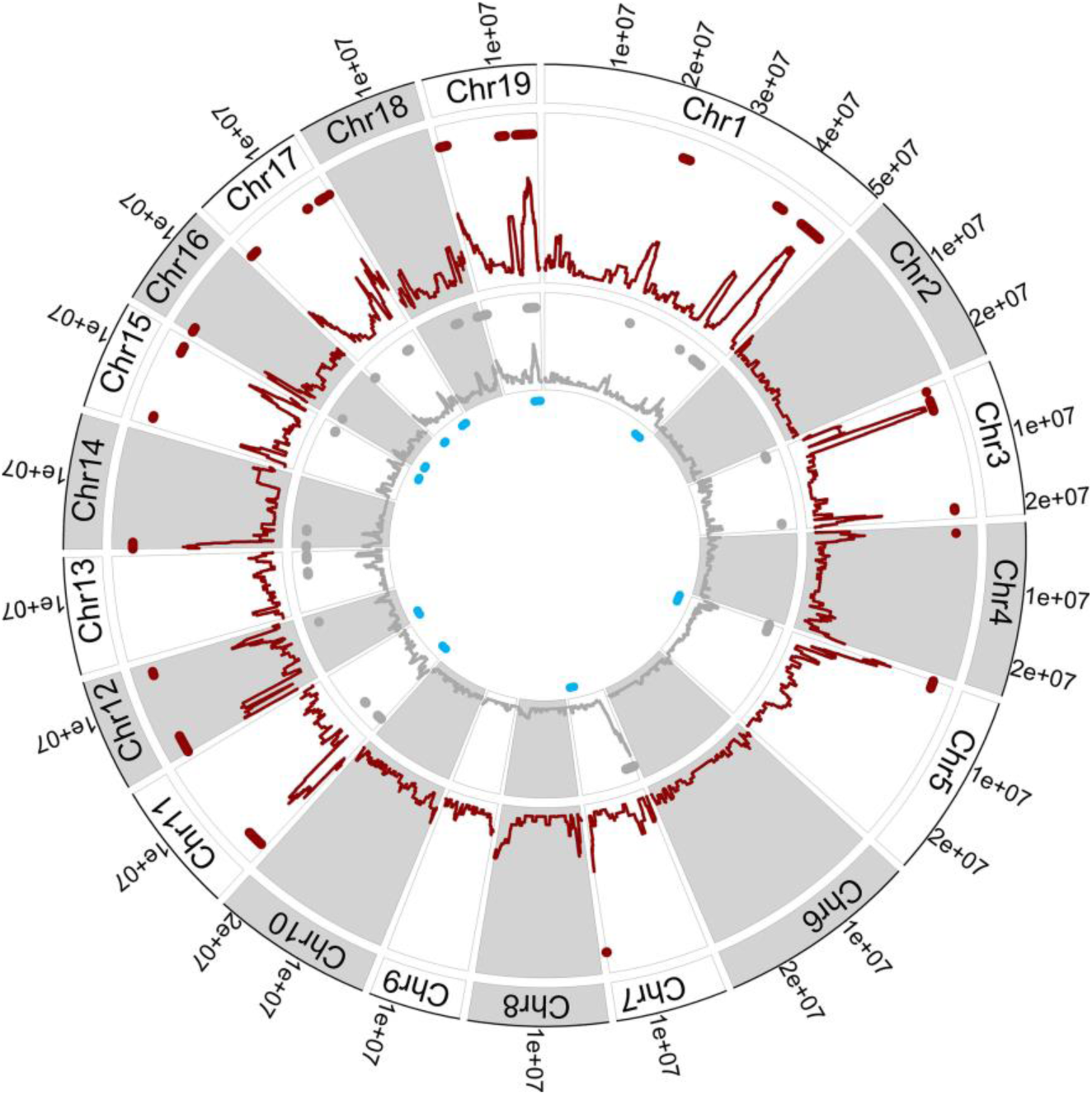
Depiction of gene CNVs and gene CNV hotspots in *Populus balsamifera*. Outer red track: all gene CNV density distribution with hotspots (red dots), inner gray track: rare gene CNV density distribution with hotspots (gray dots), inner blue dots: within species gene CNV hotspots found in *Populus nigra* and *Populus deltoides* (Pinosio et al., 2016). Figure taken from Prunier et al., 2019 (Licensed Content Author Ilga Porth, John MacKay, Nathalie Isabel, et al., License Number 5435551250896, License date Nov 24, 2022, John Wiley and Sons).

Zhang and colleagues (2019) inferred the poplar pangenome across five different sections of *Populus*, represented by poplar species native to China (sect. Tacamahaca: *P. cathayana, P. ussuriensis, P. maximowiczii, P. simonii*; sect. Leucoides: *P. lasiocarpa*; sect. Aigeiros: *P. nigra*; sect. Leuce: *P. davidiana, P. alba*; sect. Turanga: *P. euphratica*). They generated Illumina HiSeq DNA short reads, subsequently aligned them to the *P. trichocarpa* genome assembly v.3.0 and called SVs as deletions, insertions, inversions, intra- or inter-chromosomal translocations. Among the studied poplar species native to China, the number range for detected SVs was wide and ranged from 8,997 SVs in *P. euphratica* to 27,183 SVs in *P. maximowiczii*, thus for the latter, the number of SVs was three times higher. Deletions were highest in absolute numbers among SV types and highest in relative number in species *P. davidiana* (77%) and *P. alba* (76%), with numbers well above the species’ average of 69%. *P. euphratica* had a relatively high number of insertions among all SVs (19%, which is almost twice the average of all poplars studied), whereas not a single insertion was detected in *P. lasiocarpa*. The number of interchromosomal translocations exceeded that of intrachromosomal translocations and was second in relative terms (14%) to deletions (69%) among SV types, and interestingly, a high relative interchromosomal translocations number (20%) was found in *P. lasiocarpa*. Inversions were the least common SV type overall. A large number of SVs were located in coding sequences, with deletions being the most common. As the authors noted, poplars appear to be more tolerant of deletions than insertions in genic regions, probably due to the whole genome duplication events during poplar evolution and related compensatory effects. Because SVs were found predominantly in gene regions, these SVs may play a role in regulating the expression of the genes in which they were found. Normally, genes containing SVs are weakly expressed in poplars. Large chromosomal deletions were found where genes from resistance-associated gene families were clustered. The authors found significant correlations between small variants (SNPs, indels) and the density of SVs, which may be related to higher nucleotide diversity near indels and SVs, indicating certain chromosomal regions that are hotspots for genomic variation (Zhang et al., 2019).

A recent study of the population genetics of *Populus tremuloides* (quaking aspen), a widely distributed keystone species in North America, indicated that the distribution of genetic diversity, estimated at the genome level for a large number of aspen populations, is highly geographically structured, with signatures of genetic adaptation to regional climate and particularly in relation to water stress (Goessen et al., 2022). This prompted us to begin studying aspen individuals that are geographically far apart to find structural genomic features that vary between these genomes. Thus, we first looked broadly at SVs affecting this keystone species (**Figure 2.2**).

**Figure 2.2.**
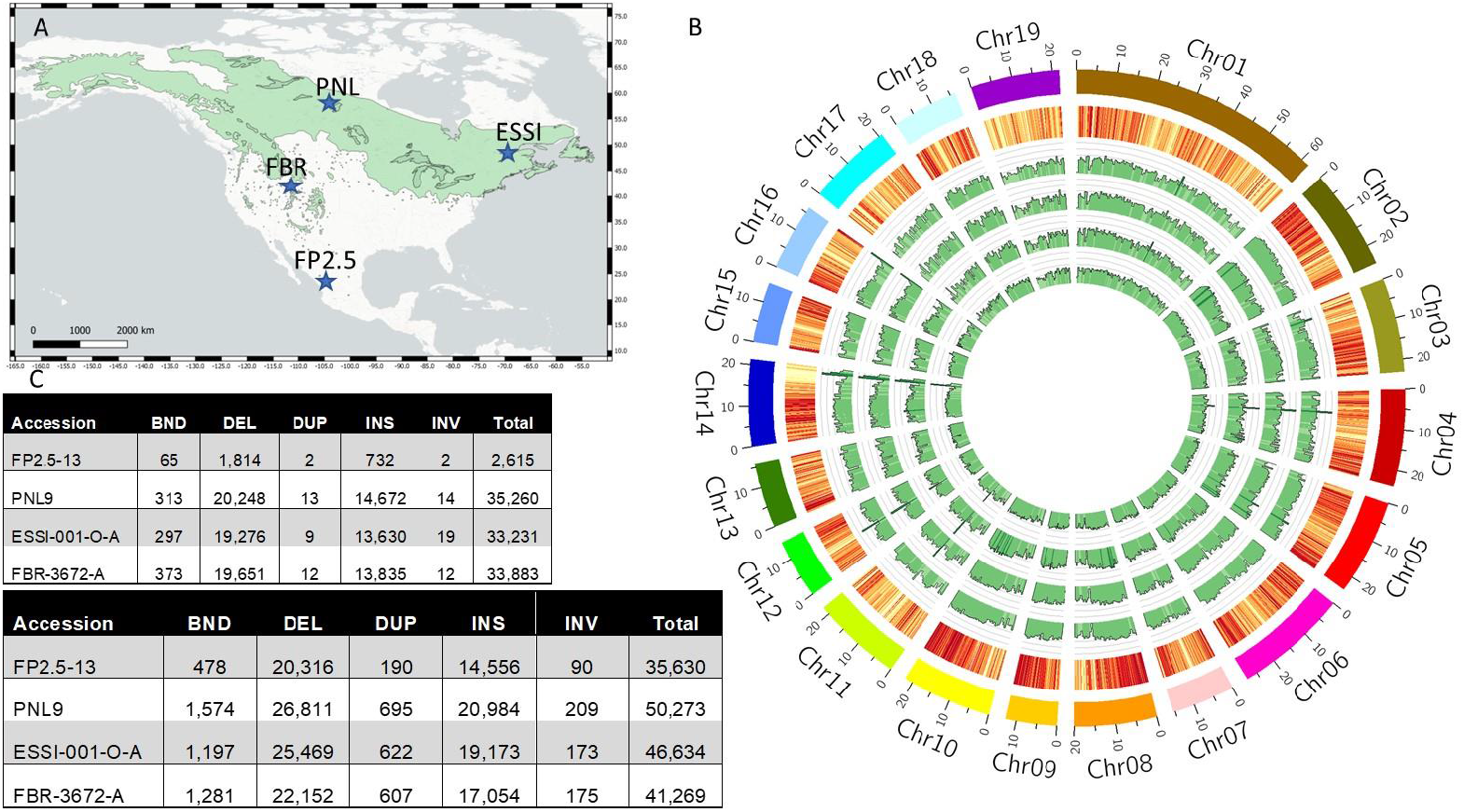
Structural variations determined across 19 chromosomes of *Populus tremuloides*. **A**. Left inset above shows *P. tremuloides*’ natural distribution across North America and depicts the four locations (blue dots) where individual samples were taken [ESSI-001-O-A (Quebec; male; 48.342 lat; -69.398 long; 39m elevation); PNL-9 (Saskatchewan; male; 58.225 lat; -103.930 long; 475m elevation); FBR-3672-A (Utah, male; 41.945 lat; -111.577 long; 2084m elevation); FP2.5-13 (Durango, male; 23.525 lat; -104.684 long; 3002m elevation)]. **B**. To the right: Track order from innermost to outermost: read coverage (light green: low, dark green: high) for: FP2.5-13, PNL9, ESSI-001-O-A, and FBR-3672-A. Gene density (yellow: low, dark red: high) in the FP2.5-13 reference genome. **C**. ONT® data from each sample were aligned against the *P. tremuloides* primary genome sequence of FP2.5-13 using NGMLR (v0.2.7) with default parameters. The alignment BAM files were analyzed by Sniffle2 (v2.0.2) to identify structural variations with precise breakpoints and minimum number of reads supporting the SV >=20 (Sedlazeck et al., 2018b). The output SV classes were defined as BND, indeterminate “break-end”; DEL, deletion; DUP, duplication; INS, insertion; and INV, inversion. The breakdown of structural variations identified in samples are summarized in the two tables: above: homozygous variant calls against *P. tremuloides* primary sequence with precise breakpoints and minimum 20 supporting reads; below: heterozygous variant calls against *P. tremuloides* primary sequence with precise breakpoints and minimum 20 supporting reads. Circos plot and SV summary were provided by Kevin Koh and Andrew Sharpe (OPAL, GIFS, Saskatchewan, Canada). I. Porth acknowledges funds under the Fonds de recherche du Québec—Nature et technologies (FRQNT) ‘Programme bilatéral de recherche collaborative Québec-Mexique’ Project 2018-265002 for whole genome sequencing and bioinformatics.

Our own preliminary results on aspen showed, again, deletions were most prevalent among SVs. The number of duplications and inversions where low amongst all five types of SVs. However, interestingly, overall, the number of insertions was relatively high, and reached proportions of on average 40%. These novel findings in aspen are very different from other poplars, where proportions of insertions hardly reached above 15%, or for certain poplar species, insertions were even completely absent (Zhang et al., 2019). In aspen, the high genetic diversity in the form of SVs suggests some rapidly evolving chromosomal regions in the genome of this species are hotspots of genetic innovation, particularly in genic regions (**Figure 2.2**) that may also be responsible for significant phenotypic variation and may be related to the species’ adaptation to local constraints. SVs could also be favoured by clonality, *i*.*e*., while they may not always be inherited sexually, they can persist in clones.

## 2.3 Induced SVs in poplar

Henry and colleagues presented an interesting concept of creating an induced structural variation system for gene dosage analysis involving gene copy numbers and functional genomics in poplar (Henry et al., 2015). This SV system was achieved by pollinating *Populus deltoides* females with a *P. nigra* male whose pollen had previously been irradiated from a γ-source (irr.). The resultant full-sib progeny encompassing hundreds of seedlings showed indels in many cases (more than half of the descendants showed them), and collectively, these chromosomal lesions covered almost all poplar genome (and on average 10 indels affecting a gene), making this resource of SVs an excellent system to provide novel insights into the effects of gene dosage on quantitative traits expression in trees. Thus, changing the gene dosage has further implications, such as on the enzyme quantity produced for example, and on the relationships of multi-component interactions (Birchler and Veitia 2021). Henry et al. (2015) used the normalized relative coverage values from Illumina short reads sequencing to identify copy numbers (dosage: normal, insertion, or deletion) in their collection of F1 individuals, *i*.*e*., in the first filial generation offspring from the cross-mating. Dosage was produced in both directions, with three-quarter deletions and one-quarter insertions. Their location was mostly at the terminal ends of chromosomes (65%). The authors confirmed that basically all indels originated from the male *P. nigra* parent. Ploidy was determined based on the parental SNP ratio in the offspring, confirming around 7% triploids for their F1 collection. Similarly, within 10 out of 15 different *Populus trichocarpa* x *P. deltoides* progeny produced by controlled crosses with various *P. trichocarpa* females, Bradshaw and Stettler (1993) detected some tri/aneuploid F1 hybrids that exhibited either transgressive phenotypes or traits predominantly similar to those of the respective female *P. trichocarpa* parents. The phenotypic variability in the *P. deltoides* x *P. nigra* (irr.) progeny produced by Henry et al. (2015) was considerable, even at a young age of the F1 seedlings. Because the insertions and deletions resulted in SVs and associated gene dosage variations in the affected genomic regions, linkage of dosage variations to phenotypic variations is possible. In this way, the genetic architecture of phenotypic traits can be dissected, and dosage-specific QTLs (dQTLs) identified. Recent research work has demonstrated the strong link between dosage effects and phenotypes for a multitude of traits (Bastiaanse et al., 2019; Rodriguez-Zaccaro et al., 2021; Bastiaanse et al., 2021), which will be further presented and discussed in the subsequent Chapter sections. A closer look at indel lines also revealed two chromoanagenetic lines (Guo et al., 2021). We understand chromoanagenesis as the highly localized restructuring (“rebirth”) of chromosomes. Plants are known to be tolerant to genome restructuring, in humans it is associated with cancer (Guo et al., 2023). The specific indel lines in poplar exhibited single chromosome fragmentation and restructuring patterns characterized by a high frequency of clustered CNVs on a single chromosome for each of the two lines (chromosome 1 and chromosome 2, respectively). These clustered dosage variations were all caused by the irradiated *P. nigra* pollen. The extreme genomic rearrangements in the 2 lines exhibited intra-chromosomal DNA junctions, and these novel junctions covered mostly genic regions, including the formation of gene-to-gene fusions. With the persistence of these two lines, there is the possibility of producing new chimeric proteins or creating new gene functions in poplar (Guo et al., 2021).

## 2.4 Alterations of phenotypes

### 2.4.1 Adaptive phenotypes associated with naturally occurring SVs in poplar

To date, a single study has examined adaptive variation in natural poplar accessions in relation to SVs. Prunier et al. (2019) examined 10 adaptive traits (six ecophysiological and four phenology traits) in controlled crosses of two balsam poplar provenances that covered a large geographical distance. Regression analysis performed between each gene CNV and each phenological and ecophysiological trait revealed a total of 23 significant associations of relative copy numbers with adaptive traits, of which 8 were associations with physiology traits, and the rest with phenology, and 13 poplar chromosomes were affected. In addition, multiple gene CNVs were found to be associated with a given adaptive trait, but no gene CNV was associated with multiple traits. Ultimately, the reported trait associations involved four phenological traits in fall and spring (timing of terminal bud initiation in fall; timing of greening of bud tip; timing of bud emergence in spring; timing of leaf emergence) and two physiological traits (photosynthetic assimilation rate; intrinsic water use efficiency). A summary of the involved genes in CNV can be found in **Table 2.1**, but their annotations suggest that they are involved in climate adaptation and/or defense against pathogens. Their effect on phenotypic variation may also be seen by the changes in their gene expression profile caused by external stimuli. This can be a topic for further study in poplars.

**Table 2.1:** List of selected poplar gene associations with leaf functional traits identified in most recent SV genomics studies in poplars. Associations are sorted by chromosomal locations.

| Association (Poplar Gene) | Gene Annotation | Putative Trait Association | SV Detection Method |
| --- | --- | --- | --- |
| *Potri.001G063700* | *AT5G49290.1* (ATRLP56, RLP56) Receptor-like protein 56 | phenology (budset timing)<sup>a</sup> | 1 |
| *Potri.001G190600* | *AT4G12760.1* replication interacting protein A | phenology (bud flush timing) | 1 |
| *Potri.001G222900* | *AT2G44480.1* (BGLU17) Beta glucosidase 17 | phenology (bud flush timing)<sup>a</sup> | 1 |
| *Potri.001G223200* | *AT5G24550.1* (BGLU32) Beta glucosidase 32 | phenology (bud flush timing)<sup>a</sup> | 1 |
| *Potri.001G399600* | *AT3G15354.1* (SPA3) SPA1-related 3 | morphology (leaf shape) | 2 |
| *Potri.001G400900* | *AT1G53000.1* (KDSB) Nucleotide-diphospho-sugar transferases superfamily protein | morphology (leaf shape) | 2 |
| *Potri.001G410300* | *AT5G54530.1* Protein of unknown function, DUF538 | morphology (leaf shape) | 2 |
| *Potri.001G415700* | *AT1G51060.1* (HTA10) histone H2A 10 | morphology (leaf shape) | 2 |
| *Potri.001G421200* | *AT4G27060.1* (CN, SPR2, TOR1) ARM repeat superfamily protein | morphology (leaf shape) | 2 |
| *Potri.001G427200* | uncharacterized | phenology (timing of green coloration of the bud tip)<sup>b</sup> | 1 |
| *Potri.001G436300* | *AT1G17220.1* (FUG1) Translation initiation factor 2, small GTP-binding protein | morphology (leaf shape) | 2 |
| *Potri.001G439000* | *AT1G16590.1* (ATREV7, REV7) DNA-binding HORMA family protein | morphology (serration; leaf shape) | 2 |
| *Potri.001G442400* | *AT4G20380.2* (LSD1) LSD1 zinc finger family protein | morphology (serration) | 2 |
| *Potri.001G448300* | *AT4G10630.1* Glutaredoxin family protein | morphology (serration; leaf shape) | 2 |
| *Potri.001G450600* | *AT1G32960.1* (ATSBT3.3, SBT3.3) Subtilase family protein | morphology (serration; leaf shape) | 2 |
| *Potri.001G453500* | *AT5G15940.1* NAD(P)-binding Rossmann-fold superfamily protein | morphology (serration; leaf shape) | 2 |
| *Potri.001G454300* | *AT1G33170.1* S-adenosyl-L-methionine-dependent methyltransferases superfamily protein | morphology (serration; leaf shape) | 2 |
| *Potri.001G454500* | *AT1G76890.2* (AT-GT2, GT2) Duplicated homeodomain-like superfamily protein | morphology (serration; leaf shape; leaf size) | 2 |
| *Potri.001G460300* | *AT3G25690.1* (CHUP1) Hydroxyproline-rich glycoprotein family protein | morphology (serration; leaf size; leaf shape) | 2 |
| *Potri.001G463200* | *AT3G26710.1* (CCB1) cofactor assembly of complex C | morphology (leaf size) | 2 |
| *Potri.001G465800* | *AT4G20940.1* Leucine-rich receptor-like protein kinase family protein | morphology (serration; leaf shape; leaf size) | 2 |
| *Potri.001G467600* | *AT3G44730.1* (ATKP1, KP1) kinesin-like protein 1 | morphology (leaf shape) | 2 |
| *Potri.001G468300* | *AT4G26500.1* (ATSUFE, CPSUFE, EMB1374, SUFE1) chloroplast sulfur E | morphology (serration; leaf size) | 2 |
| *Potri.001G468700* | *AT1G55760.1* BTB_POZ domain-containing protein | morphology (serration; leaf size) | 2 |
| *Potri.001G469800* | *AT1G55730.1* (ATCAX5, CAX5) cation exchanger 5 | morphology (serration; leaf size) | 2 |
| *Potri.001G470300* | *AT2G04690.1* Pyridoxamine 5-phosphate oxidase family protein | morphology (serration; leaf size) | 2 |
| *Potri.001G471400* | *AT5G56180.1* (ARP8, ATARP8) actin-related protein 8 | morphology (leaf size) | 2 |
| *Potri.001G471700* | *AT5G56180.1* (ARP8, ATARP8) actin-related protein 8 | morphology (serration; leaf size) | 2 |
| *Potri.002G058300* | *AT1G67930.1* Golgi transport complex protein-related | morphology (leaf shape) | 2 |
| *Potri.002G061000* | *AT1G71810.1* Protein kinase superfamily protein | morphology (serration; leaf size; leaf shape) | 2 |
| *Potri.002G063400* | *AT1G71720.1* Nucleic acid-binding proteins superfamily | morphology (serration) | 2 |
| *Potri.002G074500* | *AT1G43900.1* Protein phosphatase 2C family protein | morphology (serration; leaf size; leaf shape) | 2 |
| *Potri.002G078700* | *AT1G21640.1* (ATNADK2, NADK2) NAD kinase 2 | morphology (serration; leaf size; leaf shape) | 2 |
| *Potri.002G081200* | *AT1G77470.1* (RFC3, RFC5) replication factor C subunit 3 | morphology (serration; leaf size; leaf shape) | 2 |
| *Potri.002G082800* | *AT1G21780.1* BTB_POZ domain-containing protein | morphology (serration; leaf size) | 2 |
| *Potri.002G083500* | *AT1G44575.1* (NPQ4, PSBS) Chlorophyll A-B binding family protein | morphology (serration; leaf size; leaf shape) | 2 |
| *Potri.002G165700* | *AT5G24090.1* (ATCHIA, CHIA) Chitinase A | phenology (timing of leaf emergence) | 1 |
| *Potri.002G234200* | *AT1G60690.1* NAD(P)-linked oxidoreductase | physiology (intrinsic water-use efficiency) | 1 |
| *Potri.004G110200* | *AT1G26560.1* (BGLU40) Beta glucosidase 40 | phenology (bud flush timing)<sup>b</sup> | 1 |
| *Potri.005G001800* | *AT1G09530.1* (PAP3, PIF3, POC1) phytochrome interacting factor 3 | morphology (leaf shape) | 2 |
| *Potri.005G006500* | *AT1G09390.1* GDSL-like Lipase acylhydrolase superfamily protein | morphology (leaf shape) | 2 |
| *Potri.005G010266* | uncharacterized | morphology (leaf shape) | 2 |
| *Potri.005G012900* | *AT5G26820.1* (ATIREG3, IREG3, MAR1, RTS3) iron-regulated protein 3 | morphology (leaf shape) | 2 |
| *Potri.005G013100* | *AT2G25470.1* (AtRLP21, RLP21) Receptor-like protein 21 | phenology (timing of leaf emergence)<sup>a</sup> | 1 |
| *Potri.005G031400* | uncharacterized | phenology (timing of leaf emergence) | 1 |
| *Potri.005G067700* | *AT5G09260.1* (VPS20.2) Vacuolar protein sorting-associated | phenology (timing of leaf emergence) | 1 |
| *Potri.005G228100* | *AT4G36550.1* ARM repeat superfamily protein | morphology (serration; leaf shape) | 2 |
| *Potri.005G254100* | *AT1G42970.1* (GAPB) glyceraldehyde-3-phosphate dehydrogenase B subunit | morphology (leaf shape) | 2 |
| *Potri.005G255500* | *AT1G42550.1* (PMI1) plastid movement impaired1 | morphology (serration) | 2 |
| *Potri.006G005300* | *AT5G28300.1* Duplicated homeodomain-like protein | physiology (photosynthetic assimilation rate) | 1 |
| *Potri.009G089200* | *AT3G19590.1* (BUB3.1) Transducin_WD40 repeat-like superfamily protein | morphology (serration) | 2 |
| *Potri.009G089500* | *AT2G14820.1* (NPY2) Phototropic-responsive NPH3 family protein | morphology (serration) | 2 |
| *Potri.009G092700* | *AT2G18990.1* (TXND9) thioredoxin domain-containing protein 9 homolog | morphology (serration) | 2 |
| *Potri.009G092800* | *AT2G14910.1* uncharacterised protein | morphology (serration) | 2 |
| *Potri.009G094400* | *AT5G44750.1* (ATREV1, REV1) DNA-directed DNA polymerases | morphology (serration; leaf size) | 2 |
| *Potri.009G097900* | *AT4G34150.1* Calcium-dependent lipid-binding (CaLB domain) family protein | morphology (leaf size; leaf shape) | 2 |
| *Potri.009G100900* | *AT4G34020.1* Class I glutamine amidotransferase-like superfamily protein | morphology (leaf size) | 2 |
| *Potri.009G109300* | *AT2G15960.1* stress-induced protein | morphology (serration) | 2 |
| *Potri.009G111700* | *AT1G49380.1* cytochrome c biogenesis protein family | morphology (serration) | 2 |
| *Potri.009G113200* | *AT1G47740.1* PPPDE putative thiol peptidase family protein | morphology (serration) | 2 |
| *Potri.009G115500* | *AT4G34490.1* (ATCAP1, CAP 1, CAP1) cyclase associated protein 1 | morphology (serration) | 2 |
| *Potri.009G122300* | *AT2G21370.1* (XK-1, XK1) xylulose kinase-1 | morphology (serration) | 2 |
| *Potri.009G124800* | *AT2G21280.1* (ATSULA, GC1, SULA) NAD(P)-binding Rossmann-fold superfamily protein | morphology (serration) | 2 |
| *Potri.009G132500* | *AT4G38660.1* Pathogenesis-related thaumatin superfamily protein | morphology (serration; leaf shape) | 2 |
| Potri.009G134900 | AT4G35040.1 (bZIP19) Basic-leucine zipper (bZIP) transcription factor family protein | morphology (leaf shape) | 2 |
| Potri.009G137500 | AT2G19090.1 Protein of unknown function (DUF630 and DUF632) | morphology (serration; leaf shape) | 2 |
| Potri.009G138500 | AT1G31830.1 Amino acid permease family protein | physiology (intrinsic water-use efficiency) | 1 |
| Potri.009G159000 | AT2G27090.1 Protein of unknown function (DUF630 and DUF632) | morphology (serration; leaf shape) | 2 |
| Potri.009G160100 | AT2G27350.1 OTU-like cysteine protease family protein | morphology (serration) | 2 |
| Potri.009G161800 | AT2G27290.1 Protein of unknown function (DUF1279) | morphology (serration) | 2 |
| Potri.009G164500 | AT3G44460.1 (AtbZIP67, DPBF2) Basic-leucine zipper (bZIP) transcription factor family protein | morphology (serration; leaf shape) | 2 |
| Potri.009G165300 | AT2G17980.1 (ATSLY1) Sec1_munc18-like (SM) proteins superfamily | morphology (serration) | 2 |
| Potri.009G169500 | AT4G38210.1 (ATEXP20, ATEXPA20, ATHEXP ALPHA 1.23, EXP20, EXPA20) expansin A20 | morphology (serration; leaf shape) | 2 |
| Potri.010G171600 | AT1G79550.1 (PGK) phosphoglycerate kinase | morphology (leaf shape) | 2 |
| Potri.010G173000 | AT1G79510.1 Uncharacterized conserved protein (DUF2358) | morphology (leaf shape) | 2 |
| Potri.011G027900 | AT4G23180.1 (CRK10, RLK4) Cysteine-rich RECEPTOR-like protein kinase | physiology (intrinsic water-use efficiency) | 1 |
| Potri.011G154800 | AT4G20380.2 (LSD1) LSD1 zinc finger family protein | morphology (leaf shape) | 2 |
| Potri.011G156900 | AT1G73720.1 (SMU1) transducin family protein _ WD-40 repeat family protein | morphology (leaf shape) | 2 |
| Potri.011G157000 | AT4G14350.1 AGC (cAMP-dependent, cGMP-dependent and protein kinase C) kinase family protein | morphology (leaf shape) | 2 |
| Potri.012G126000 | AT5G62390.1 (ATBAG7, BAG7) BCL-2-associated athanogene 7 | morphology (leaf shape) | 2 |
| Potri.012G126400 | AT4G25240.1 (SKS1) SKU5 similar 1 | morphology (leaf shape) | 2 |
| Potri.012G131500 | AT5G51750.1 (ATSBT1.3, SBT1.3) subtilase 1.3 | morphology (leaf shape) | 2 |
| Potri.013G007800 | AT1G56500.1 haloacid dehalogenase-like hydrolase family protein | morphology (serration) | 2 |
| Potri.013G008500 | AT5G26850.1 uncharacterized protein | morphology (serration) | 2 |
| Potri.013G012200 | AT5G27930.1 Protein phosphatase 2C family protein | morphology (serration) | 2 |
| Potri.013G017166 | uncharacterised | morphology (serration) | 2 |
| Potri.013G020800 | AT5G27560.1 Domain of unknown function (DUF1995) | morphology (serration) | 2 |
| *Potri.013G021500* | *AT1G54990.1* (AXR4, RGR, RGR1) alpha, beta-Hydrolases superfamily protein | morphology (serration) | <sup>2</sup> |
| *Potri.014G007200* | *AT2G22840.1* (AtGRF1, GRF1) growth-regulating factor 1 | morphology (serration; leaf shape; leaf size) | <sup>2</sup> |
| *Potri.014G008100* | *AT4G37750.1* (ANT, CKC, CKC1, DRG) Integrase-type DNA-binding superfamily protein | morphology (serration; leaf shape) | <sup>2</sup> |
| *Potri.014G012800* | *AT2G22840.1* (AtGRF1, GRF1) growth-regulating factor 1 | morphology (serration; leaf shape; leaf size) | <sup>2</sup> |
| *Potri.014G014300* | *AT3G49810.1* ARM repeat superfamily protein | morphology (serration; leaf shape; leaf size) | <sup>2</sup> |
| *Potri.014G016000* | *AT1G27460.1* (NPGR1) no pollen germination related 1 | morphology (serration; leaf shape; leaf size) | <sup>2</sup> |
| *Potri.014G017900* | *AT2G26680.1* FkbM family methyltransferase | morphology (leaf shape) | <sup>2</sup> |
| *Potri.014G020200* | *AT5G43100.1* Eukaryotic aspartyl protease family protein | morphology (serration; leaf size) | <sup>2</sup> |
| *Potri.014G021900* | *AT1G47200.1* (WPP2) WPP domain protein 2 | morphology (leaf size) | <sup>2</sup> |
| *Potri.014G023400* | *AT3G46630.1* Protein of unknown function (DUF3223) | morphology (serration) | <sup>2</sup> |
| *Potri.014G023400* | *AT3G46630.1* Protein of unknown function (DUF3223) | morphology (serration; leaf shape) | <sup>2</sup> |
| *Potri.014G025700* | *AT4G27410.2* (ANAC072, RD26) NAC (No Apical Meristem) domain transcriptional regulator superfamily protein | morphology (serration; leaf shape; leaf size) | <sup>2</sup> |
| *Potri.014G032400* | *AT1G47420.1* (SDH5) succinate dehydrogenase 5 | morphology (leaf size; leaf shape) | <sup>2</sup> |
| *Potri.014G047500* | *AT2G44830.1* Protein kinase superfamily protein | morphology (leaf shape) | <sup>2</sup> |
| *Potri.014G050300* | *AT1G02150.1* Tetratricopeptide repeat (TPR)-like superfamily protein | morphology (serration; leaf shape; leaf size) | <sup>2</sup> |
| *Potri.014G050300* | *AT1G02150.1* Tetratricopeptide repeat (TPR)-like superfamily protein | morphology (serration; leaf shape; leaf size) | <sup>2</sup> |
| *Potri.014G053600* | *AT3G60220.1* (ATL4, TL4) TOXICOS EN LEVADURA 4 | morphology (serration; leaf shape; leaf size) | <sup>2</sup> |
| *Potri.014G055800* | *AT2G44950.1* (HUB1, RDO4) histone mono-ubiquitination 1 | morphology (leaf shape) | <sup>2</sup> |
| *Potri.014G060100* | *AT2G45150.1* (CDS4) cytidine diphosphate diacylglycerol synthase 4 | morphology (serration; leaf shape; leaf size) | <sup>2</sup> |
| *Potri.014G066700* | *AT2G45190.1* (AFO, FIL, YAB1) Plant-specific transcription factor YABBY family protein | morphology (leaf shape) | <sup>2</sup> |
| *Potri.014G068700* | *AT2G45340.1* Leucine-rich repeat protein kinase family protein | morphology (serration) | <sup>2</sup> |
| *Potri.014G070000* | *AT4G12080.1* (AHL1, ATAHL1) AT-hook motif nuclear-localized protein 1 | morphology (serration; leaf shape; leaf size) | <sup>2</sup> |
| *Potri.014G070100* | *AT5G51600.1* (ATMAP65-3, MAP65-3, PLE) Microtubule associated protein (MAP65_ASE1) family protein | morphology (serration; leaf shape; leaf size) | <sup>2</sup> |
| *Potri.014G071800* | *AT2G45480.1* (AtGRF9, GRF9) growth-regulating factor 9 | morphology (serration; leaf shape; leaf size) | <sup>2</sup> |
| *Potri.014G071900* | *AT2G45500.1* AAA-type ATPase family protein | morphology (serration; leaf size; leaf shape) | <sup>2</sup> |
| *Potri.014G072800* | *AT3G61050.1* (NTMC2T4, NTMC2TYPE4) Calcium-dependent lipid-binding (CaLB domain) family protein | morphology (serration; leaf shape; leaf size) | <sup>2</sup> |
| *Potri.014G072900* | *AT2G45590.1* Protein kinase superfamily protein | morphology (serration; leaf shape; leaf size) | <sup>2</sup> |
| *Potri.014G075600* | *AT4G00752.1* UBX domain-containing protein | morphology (serration; leaf shape; leaf size) | <sup>2</sup> |
| *Potri.014G076500* | *AT2G45720.1* ARM repeat superfamily protein | morphology (serration; leaf shape; leaf size) | <sup>2</sup> |
| *Potri.014G077800* | *AT5G51980.1* Transducin_WD40 repeat-like superfamily protein | morphology (leaf shape) | <sup>2</sup> |
| *Potri.014G078100* | *AT1G01910.1* P-loop containing nucleoside triphosphate hydrolases superfamily protein | morphology (serration; leaf size; leaf shape) | <sup>2</sup> |
| *Potri.014G080600* | *AT4G00590.1* N-terminal nucleophile aminohydrolases (Ntn hydrolases) superfamily protein | morphology (serration; leaf shape; leaf size) | <sup>2</sup> |
| *Potri.014G083400* | *AT5G60230.2* (ATSEN2, SEN2) splicing endonuclease 2 | morphology (leaf shape; leaf size) | <sup>2</sup> |
| *Potri.014G084200* | *AT4G00460.2* (ATROPGEF3, ROPGEF3) RHO guanyl-nucleotide exchange factor 3 | morphology (serration; leaf shape; leaf size) | <sup>2</sup> |
| *Potri.014G084900* | *AT2G45910.1* U-box domain-containing protein kinase family protein | morphology (serration; leaf shape; leaf size) | <sup>2</sup> |
| *Potri.014G085700* | *AT4G00370.1* (ANTR2, PHT4;4) Major facilitator superfamily protein | morphology (serration) | <sup>2</sup> |
| *Potri.014G091200* | *AT2G46170.1* Reticulon family protein | morphology (serration; leaf size; leaf shape) | <sup>2</sup> |
| *Potri.014G093900* | *AT4G01150.1* similar to CALMODULIN 2, poplar CaM939 calmodulin | morphology (serration; leaf shape; leaf size) | <sup>2</sup> |
| Potri.014G100400 | AT4G10840.1 Tetratricopeptide repeat (TPR)-like superfamily protein | morphology (serration; leaf size; leaf shape) | <sup>2</sup> |
| Potri.014G104300 | AT1G01120.1 (KCS1) 3-ketoacyl-CoA synthase 1 | morphology (serration; leaf shape; leaf size) | <sup>2</sup> |
| Potri.014G104600 | AT4G00820.1 (iqd17) IQ-domain 17 | morphology (serration; leaf shape; leaf size) | <sup>2</sup> |
| Potri.014G125600 | AT2G47490.1 (ATNDT1, NDT1) NAD<sup>+</sup> transporter 1 | morphology (serration; leaf size; leaf shape) | <sup>2</sup> |
| Potri.014G160600 | AT2G04530.1 (CPZ, TRZ2) Metallo-hydrolase oxidoreductase superfamily protein | morphology (serration) | <sup>2</sup> |
| Potri.015G018600 | AT5G38280.1 (PR5K) PR5-like receptor kinase | phenology (bud flush timing)<sup>b</sup> | <sup>1</sup> |
| Potri.016G046200 | AT3G12120.2 (FAD2) Fatty acid desaturase 2 | phenology (timing of green coloration of the bud tip) | <sup>1</sup> |
| Potri.016G118500 | AT2G29420.1 (ATGSTU7, GST25, GSTU7) Glutathione S-transferase tau 7 | phenology (budset timing) | <sup>1</sup> |
| Potri.016G127000 | AT1G45616.1 (AtRLP6, RLP6) Receptor-like protein 6 | physiology (intrinsic water-use efficiency) | <sup>1</sup> |
| Potri.017G009400 | AT1G66920.2 Protein kinase superfamily protein | physiology (intrinsic water-use efficiency) | <sup>1</sup> |
| Potri.017G024300 | AT3G60470.1 plant protein of unknown function (DUF247) | phenology (timing of green coloration of the bud tip) | <sup>1</sup> |
| Potri.018G075100 | AT3G53400.1 (CPuORF47) Conserved Peptide Upstream Open Reading Frame 47 | physiology (photosynthetic assimilation rate) | <sup>1</sup> |
| Potri.019G008100 | AT1G29670.1 GDSL-like Lipase_Acylhydrolase superfamily protein | morphology (serration; leaf size; leaf shape) | <sup>2</sup> |
| Potri.019G013100 | AT5G28640.1 (AN3, ATGIF1, GIF, GIF1) SSXT family protein | morphology (serration; leaf size; leaf shape) | <sup>2</sup> |
| Potri.019G070300 | AT5G17680.1 Disease resistance protein TIR-NBS-LRR class | phenology (timing of green coloration of the bud tip) | <sup>1</sup> |
| Potri.019G109900 | AT4G29990.1 Leucine-rich repeat transmembrane protein kinase | phenology (timing of leaf emergence) | <sup>1</sup> |

<sup>1</sup> aCGH technology screens of *P. balsamifera* F1 progeny (Prunier al., 2019)
<sup>2</sup> Illumina short reads sequencing of *P. deltoides* x *P. nigra* F1 hybrids generated by using irradiated (irr.) *P. nigra* pollen (Bastiaanse et al., 2021)
<sup>a</sup> within dQTL region(s) in *P. deltoides* x *P. nigra* (irr.) for fall phenology (Bastiaanse et al., 2019)
<sup>b</sup> within dQTL region(s) in *P. deltoides* x *P. nigra* (irr.) for spring phenology (Bastiaanse et al., 2019)

### 2.4.2 Variation in adaptive phenotypes related to artificially induced SVs in poplar

Indel lines described by Henry et al. (2015) also served as a resource to examine the effects of induced gene dosage variation on spring and fall phenology (leaf bud expansion, leaf yellowing and senescence, and duration of canopy greening) and related biomass traits (tree height, diameter, and volume) (Bastiaanse et al., 2019). Among the approximately 600 indel lines phenotyped for these traits over a four-year period, the authors found that biomass traits were almost always negatively affected, including a significant reduction in tree height. For phenology traits, significantly later bud flushing and shortened duration of green canopy was observed in the extreme indel lines. Phenotypic variation attributable to genotypic variation (trait heritability) was moderate to high for phenology and biomass traits. Moderate to high genetic correlations among traits were also observed. In addition, no difference was found in the direction of effect sizes between deletions and insertions affecting phenotypes, but length size had an additional effect for deletions but not for insertions on phenotype. Over half of the poplar genome was found sensitive to dosage variation for at least one phenotype, indicating that dosage balance on a genomic scale determines quantitative trait variation. Quantitative trait variation at specific chromosomal locations was found to be sensitive to gene dosage, leading to the discovery of dQTLs (Bastiaanse et al., 2019). The strongest impact of such loci was found in phenological traits, which also showed high repeatability across years. Both increases and decreases in relative dosage in indel-bearing lines were associated with proportional modulation of phenotype above and below that of control lines (without indel). Such loci could therefore be prime targets for tree improvement.

### 2.4.3 Variation in wood anatomy traits related to artificially induced SVs in poplar

Rodriguez-Zaccaro et al. (2021) used a subset (c.200 lines, ensuring 95% of the genome was covered by at least one indel) of the clonally replicated and genomically defined indel lines generated by the work of Henry et al. (2015) to specifically examine the effects of gene dosage on vessel traits, the proportion of trait variance explained by dQTLs. Phenotypes were assessed in a single growing season, and some vessel traits were height-corrected prior to further analyses. Indels have resulted in phenotypic variation beyond that caused by allelic segregation found in control genotypes without induced indels. In fact, gene dosage correlated with the measured quantitative traits, identifying the QTLs for those traits. Traits also showed significant and moderate to higher heritability. The proportion of anatomical trait variance explained by dQTLs ranged between 3 to 15%, depending on the trait considered. These dosage-responsive QTLs contained between 63 and 380 annotated genes, depending on the trait and QTL position. Additionally, correlated traits showed evidence of shared dQTLs, but also trait-specific dQTLs. These results also provide information on which traits can be targeted independently through breeding approaches, e.g., vessel traits independent of tree height growth (Rodriguez-Zaccaro et al., 2021).

### 2.4.4 Variation in leaf morphology related to artificially induced SVs in poplar

Finally, the indel lines created by Henry and co-workers (2015), including clonal replicates, also served Bastiaanse et al. (2021) to identify dosage sensitive co-expressed gene modules, enriched in gene dosage-specific expression QTLs, and associated with morphological features. In fact, gene dosage can modulate the expression of a gene directly (*cis*) or indirectly through a transcription factor (*trans*) that is affected by dosage variation. The authors phenotyped 600 lines for several leaf morphometric traits and produced transcriptomic data for c.28% of these indel lines. First, a classical QTL analysis was performed for morphometric traits using dosage variation in correlation analysis to assess the frequency and magnitude of the effects of dosage-responsive loci on these leaf traits; second, RNAseq data were used to perform QTL analysis for expression phenotypes that depend on dosage variation to assess the effects of dosage on the expression of individual genes affecting leaf morphology (differentially expressed genes from pools of phenotypically extreme mutant lines were compared with QTL locations). Finally, co-expressed gene modules were correlated with variation in leaf morphology traits. Integration of all these datasets and analyses resulted in a list of 116 unique candidate genes (**Table 2.1**) underlying morphological variation (Bastiaanse et al., 2021). The marked effect of induced dosage variation on leaf morphology suggests that naturally occurring SVs may also contribute to morphological changes in wild populations. Dosage variation could also cause rapid morphological changes during speciation.

## 2.5 Future directions for integrating knowledge for breeding applications

The study by Bastiaanse and colleagues (2019) provided the first insights into the importance of gene dosage variation for expression variation of adaptive phenotypes in poplars. The study by Prunier et al. (2019) was the first to reveal the extent of intraspecific variation of CNVs in poplar, and this in relation to adaptive phenology and physiology. Due to their generally faster mutation rates compared to SNPs (Prunier et al., 2019), SVs are effective drivers of evolutionary and ecological processes (Mérot et al., 2020) and enable plants (including poplars) to adapt to environmental stimuli (biotic and abiotic stressors) more effectively, as they do so more rapidly. However, it is crucial to (a) provide common standards for SV detection that would facilitate comparative studies (discordance between different technologies can be as high as 80%; Alkan et al., 2011) and (b) develop new theoretical approaches tailored to the diversity of SV types with different evolutionary trajectories (Mérot et al., 2020).

Domesticated crops such as soybean, rice, tomato, barley, or wheat were among the first plants to attract interest in SVs (cited in Hämälä et al., 2021). Recently, cocoa (*Theobroma cacao*) and barley (*Hordeum vulgare*) were evaluated for direct effects of SVs on plant fitness and crop yield (Hämälä et al., 2021; Weisweiler et al., 2022). Both studies, although using different technologies to detect SVs - haplotype-resolved genome assembly vs. short-read sequencing - showed that SVs affect transcript abundance, and some SVs even have positive effects on fitness (including pathogen resistance). While most SVs in wild accessions, as shown for cocoa, are subject to purifying selection – that is, these SVs have been selected against and are rare in the population -, several SVs in domesticated crops are associated with important traits for plant improvement, such as abiotic stress tolerance (Hämälä et al., 2021). The use of SV clusters (especially insertions) determined from short-read sequencing data in phenotypic prediction models yielded, on average, higher predictive ability than SNPs for a range of yield and plant architecture related quantitative traits in barley (Weisweiler et al., 2022). This is because (1) gene-associated SV clusters (i.e., SVs clustering within 50 bp and found within and 5 kb upstream/downstream around genes) are more likely to underlie causal genes than SNPs, a significant proportion of SVs (c.10%) is directly associated with gene-specific gene expression, and also because (3) SV clusters have less linkage disequilibrium between them, thus, collectively, making SV clusters more informative than SNPs for phenotypic prediction and genomic selection. DNA SVs may thus help to explain the missing heritability of quantitative traits. In fact, Della Coletta and colleagues (2021) specifically proposed the integration of information on genome content variation for crop improvement, in part due to the growing amount of pangenome information for plant species. Hence, SVs may become potent new DNA marker types that provide enhanced ability to select individuals for complex quantitative traits of interest (Lorenz et al., 2011) by estimating SV effects across the genome. While “characterizing the SV landscape at the breeding program level” (Della Coletta et al., 2021) needs to take into account different approaches to genome assembly (including graph-based pangenomics algorithms; Golicz et al., 2015), consideration of the SV-based pangenome promises a paradigm shift in plant improvement (Della Coletta et al., 2021).

In poplars, breeding strategies are mostly aimed at exploiting hybrid vigor where the resulting heterozygous F1 progeny has led to superior characteristics compared to either parent producing these hybrid crosses. Heterosis in plants can be due to dominance, overdominance, and epistasis (Springer and Stupar 2007), and is usually restricted to the F1 generation; it is noteworthy that not all crosses produce transgressive phenotypes (Wang et al., 2015). Understanding the molecular mechanisms leading to transgressive phenotypes is key to selecting molecular markers in support of hybrid breeding endeavours. Detection of structural heterozygosity in conjunction with correlated methylation and transcription patterns would allow comparisons of SV effects among individuals, which appears to be the basis for significant phenotypic implications including heterosis (see cited articles on poplar in this Chapter), although it will be important to also assess the predictive robustness of SV effects in different environments.

## Notes

### Competing Interest Statement

The authors have declared no competing interest.

### Summary of Updates

A typo has been corrected in the Introduction.

